# Anterograde Trans-Synaptic AAV Strategies for Probing Neural Circuitry

**DOI:** 10.1101/2019.12.24.888172

**Authors:** Brian Zingg, Bo Peng, Junxiang J. Huang, Huizhong W. Tao, Li I. Zhang

**Author notes:** Correspondence should be addressed to: L.I.Z or H.W.T.

## Abstract

Elucidating the organization and function of neural circuits is greatly facilitated by viral tools that spread transsynaptically. Adeno-associated virus (AAV) has been shown to exhibit anterograde transneuronal spread. However, the synaptic specificity of the spread and its broad application in various neural circuits remain to be explored. Here, using anatomical, functional, and molecular approaches, we provide strong evidence for the specifically preferential spread of AAV1 to post-synaptically connected neurons. Besides glutamatergic synapses made onto excitatory and inhibitory neurons, AAV1 also transsynaptically spreads through GABAergic synapses and effectively tags spinal cord neurons receiving long-distance projections from various brain regions, but exhibits little or no spread through neuromodulatory projections (e.g. serotonergic, cholinergic, and noradrenergic). Combined with newly designed intersectional and sparse labeling strategies, AAV1 can be utilized to categorize neurons according to their input sources, morphological and molecular identities. These properties make AAV a unique anterograde transsynaptic tool for establishing a comprehensive cell-atlas of the brain.

## Introduction

Viral tools that spread transsynaptically greatly facilitate research work to reveal the organization and function of neural circuits (Luo et al., 2018; Nassi et al., 2015). Adeno-associated virus (AAV) has recently been shown to be capable of anterograde transneuronal transport, with serotype 1 (AAV1) in particular exhibiting the greatest efficiency of spread (Zingg et al., 2017). Given its well established lack of toxicity and apparent transduction of only first-order postsynaptic neurons, AAV1 shows great promise as a tool for manipulating input-defined cell populations and mapping their outputs, comparing to other antero-transneuronal viral tools currently under development (Beier et al., 2011; Lo et al., 2011; Zeng et al., 2017; Gradinaru et al., 2010). This approach has become more and more widely used recently (e.g. Bennet et al., 2019; Castle et al., 2014b; Cembrowski et al., 2018; Yao et al., 2018; Wang et al., 2018; Huang et al., 2019; Beltramo & Scanziani, 2019; Centanni et al., 2019; Sengupta & Holmes, 2019; Trouche et al., 2019).

Previous work suggests that AAV1 is released at or near axon terminals, and transduced neurons downstream of the injection site show a high probability of receiving functional synaptic input in slice recording experiments (Zingg et al., 2017). However, the extent to which AAV1 spreads exclusively to synaptically connected neurons remains uncertain. In addition, despite clear evidence for the active trafficking of AAV-containing vesicles down the axon (Castle et al, 2014a; Castle et al., 2014b), exactly how AAV is eventually released (e.g. through synaptic or extrasynaptic vesicle fusion) remains unknown. Addressing these questions will be essential for establishing the synaptic nature of AAV transneuronal transduction.

AAV1 has been shown to efficiently transduce both excitatory and inhibitory neurons downstream of a variety of glutamatergic corticofugal pathways (Zingg et al., 2017; Yao et al., 2018; Wang et al., 2018; Centanni et al., 2019; Bennet et al., 2019). In addition, this efficiency appears to be critically dependent on viral titer, as reducing the titer from 10^13^ to 10^11^ GC/mL completely eliminates transneuronal spread (Zingg et al., 2017). Given the molecular diversity among different cell-types in the brain, it remains uncertain whether differences in cell surface receptor expression, intracellular trafficking, or synapse type might limit the efficiency of AAV spread in certain pathways. In particular, transneuronal spread through inhibitory projection neurons or neuromodulatory cell populations has yet to be directly examined. Moreover, whether or not axon length might diminish spread (e.g. from cortex to spinal cord) remains to be tested.

In this study, we systematically examine the synaptic specificity of AAV1 transneuronal transport using a variety of anatomical, functional, and molecular approaches. We find a strong correspondence between presynaptic connectivity and postsynaptic labeling for different pathways, and find that co-expression of tetanus toxin light chain, an inhibitor of presynaptic vesicle fusion, nearly abolishes transsynaptic spread of AAV. In addition, we establish that AAV1 spreads efficiently through inhibitory projection pathways, as well as long-range pathways from the brain to the spinal cord, but shows little or no spread through all neuromodulatory projections examined. Lastly, we expand the application potential for this technique by combining Flp- and Cre-dependent mapping in dual-reporter mice and incorporate its use with approaches for the sparse labeling of input- and genetically-defined neurons.

## Results

### Synaptic specificity of AAV transneuronal spread

Previously, we observed that AAV1-Cre spreads only to first-order downstream neurons in expected target regions, does not leak from axons of passage, and labels neurons that show a high probability of receiving functional presynaptic input (Zingg et al., 2017). These observations are suggestive of a transsynaptic mechanism of spread, however the extent to which AAV1 passes exclusively through synaptic connections remains uncertain (Figure 1A). To provide a deeper understanding of the synaptic nature of AAV1 transport, we systematically characterized (1) the specificity of the spread to only expected cell-types in a well-defined circuit, (2) the probability of functional connectivity between labeled and non-labeled cells in a target region, and (3) whether AAV spread is dependent on synaptic vesicle release.

**Figure 1.**
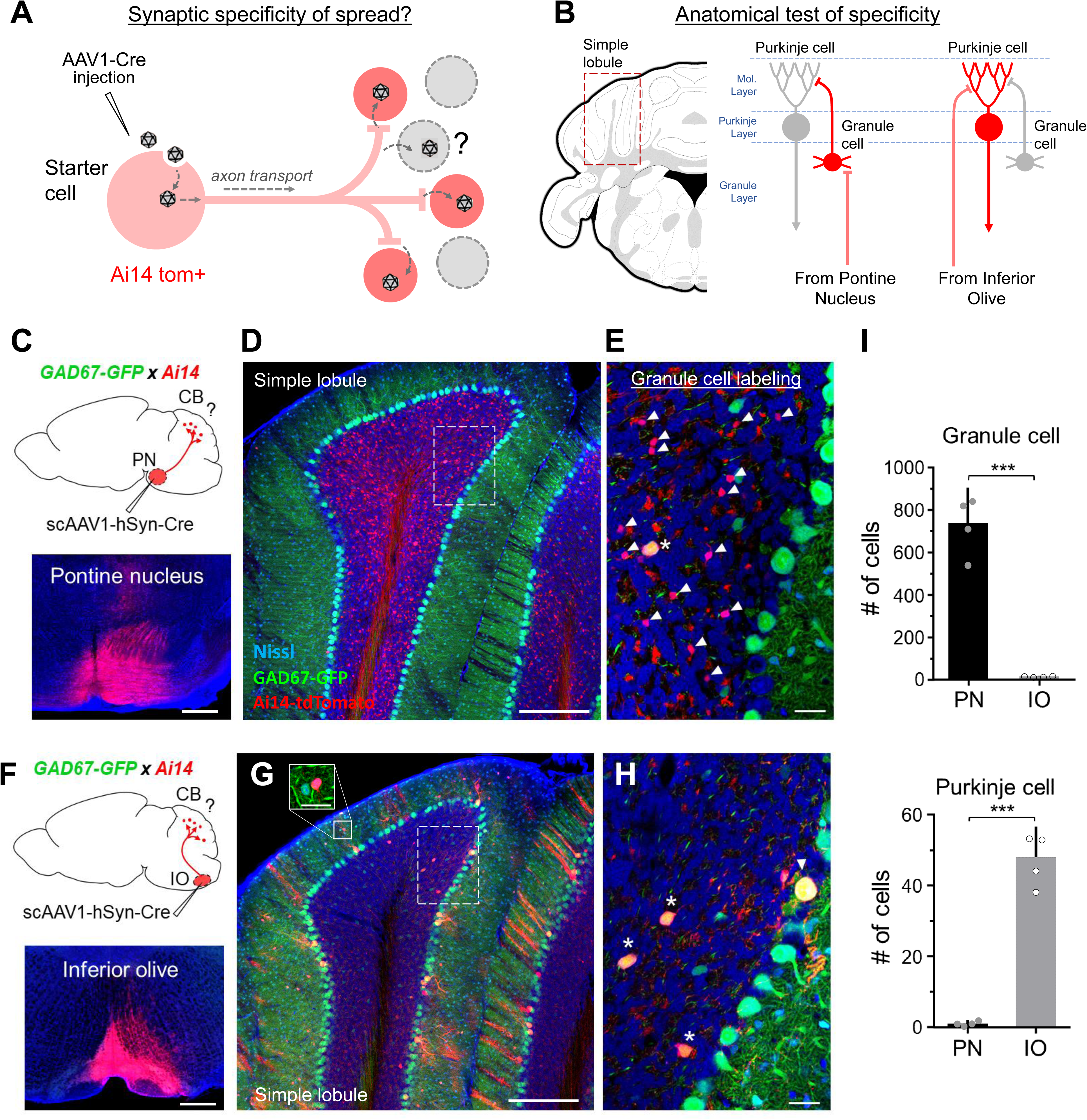
Anatomical evidence for the synaptic specificity of viral spread.

A. Illustration to show that following injection in an upstream brain region, AAV1-Cre is transported down axons and may be released through the synapse to transduce post-synaptically connected neurons (red cells, in Cre-dependent tdTomato background). The extent to which extra-synaptic release of virus may contribute to the local transduction of unconnected cell types (gray cells) remains unclear.
B. Strategy for testing the synaptic specificity of viral spread in an anatomically defined circuit. Post synaptic labeling was examined in the simple lobule of the cerebellar cortex following injections in the inferior olive or pontine nucleus (left panel). Mossy fiber afferents from the pontine nucleus are known to innervate granule cells but not Purkinje cells (right). On the other hand, climbing fiber afferents from the inferior olive pass through the granule layer and selectively innervate Purkinje cells, but not granule cells. Locations of cell bodies in different layers of the cerebellar cortex are indicated by dashed lines (Molecular layer, Purkinje layer, or Granule layer).
C. Approach for labeling neurons postsynaptic to mossy fibers in the cerebellar cortex (CB). The scAAV1-hSyn-Cre was injected into the pontine nucleus (PN) of Ai14 × GAD67-GFP transgenic mice. Bottom panel, example injection site (red). Blue, fluorescent Nissl stain. Scale bar, 500 µm.
D. A coronal section through simple lobule showing pontine afferents and postsynaptic neurons labeled in the granule layer of the cerebellum (red). GAD67-GFP+ Purkinje cells and molecular layer interneurons were labeled in green. Blue, fluorescent Nissl stain. Scale bar, 250 µm.
E. Higher magnification view of the dashed region shown in (D). tdTomato+/GFP- granule cells (red, co-stained with Nissl, blue, arrowheads) and mossy fiber terminals (red) were observed in the granule layer, along with large GAD67-GFP+ neurons (yellow, asterisk). Scale bar, 25 µm.
F. Approach for labeling neurons postsynaptic to climbing fibers in CB. The scAAV1-hSyn-Cre was injected into the inferior olive of Ai14 × GAD67-GFP mice. Bottom panel, example injection site (red). Scale bar, 500 µm.
G. Climbing fiber afferents and postsynaptic neurons (red) labeled in the granule, Purkinje, and molecular layers of the simple lobule. Most labeled neurons in the granule cell layer co-localizing with GAD67-GFP+ expression are presumed Golgi cells which can be innervated by climbing fibers. Scale bar, 250 µm. Solid box shows close up of a molecular layer interneuron (MLI). Scale bar, 25 µm.
H. Higher magnification view of the dashed region shown in (G). tdTomato+/GFP+ neurons were observed in the granule layer (yellow, asterisks), Purkinje cell layer (yellow, arrowhead), and molecular layer (cells shown in G).
I. Quantification of the total number of granule (upper) and Purkinje cells (lower) counted across 4 sections of the simple lobule for each animal injected in PN or IO (mean plotted for n = 4 animals for each pathway). Error bar = SD. ***p < 0.001, t-test.

### Anatomical examination of AAV transsynaptic spread

To provide a broad estimation of the synaptic specificity of AAV1 transneuronal spread, we took advantage of two different unidirectional pathways that converge on distinct cell-types within the cerebellar cortex (Figure 1B). Neurons in the pontine nucleus (PN) project to the granule layer of the cerebellum and form well-characterized connections with granule cells (Palay & Chan-Palay, 1974). On the other hand, neurons in the inferior olive (IO) send axons through the granule layer and terminate primarily onto the dendrites of Purkinje cells (Palay & Chan-Palay, 1974; Mathews et al., 2012). We therefore expected injections of AAV1-Cre in the PN to label granule cells, but not Purkinje cells, and injections in the IO to label Purkinje cells, but not granule cells. To test this, we first injected self-complementary (sc)AAV1-hSyn-Cre into the PN or IO of Ai14 × GAD67-GFP mice (Figure 1C, 1F). This AAV is packaged with a double-stranded DNA construct and exhibits greater transduction efficiency (McCarty, 2008) and increased anterograde transneuronal labeling when compared with similar AAVs containing single-stranded DNA (see Materials and Methods and Figure S1). GAD67-GFP expression was mainly used to facilitate identification of Purkinje cells. Following a 2-week post-injection survival time, PN injections yielded hundreds of small tdTomato+/GFP− cells in the granule layer (presumed granule cells) (Figure 1D-E), while IO injections labeled giant tdTomato+/GFP+ Purkinje cells in the Purkinje layer (Figure 1G-H). We here focus our estimation of the synaptic specificity of viral spread using Purkinje and granule cells because they are the only cell types receiving well established point-to-point synaptic contacts from PN or IO pathways (Brown et al., 2012; Galliano et al., 2013; Szapiro et al., 2007; Coddington et al., 2013). To quantify this labeling, we focused on 4 consecutive sections of the contralateral simple lobule (SIM; see Materials and Methods) and counted the total number of granule cells and Purkinje cells across all 4 sections for each animal (n = 4 animals for both PN and IO). As expected, PN injections labeled a large number of granule cells (average 735 cells per animal), while IO injections labeled a negligible number of granule cells (average 13 cells per animal) (Figure 1I, upper). Similarly, IO injections robustly labeled Purkinje cells (average 48 cells per animal), while PN injections labeled little or no Purkinje cells (average 1 cell per animal) (Figure 1I, lower). Given that each pathway is expected to innervate one cell-type and exclude the other, the non-zero values observed for unexpected cell-types may represent potential leakage of the virus to non-synaptically connected neurons. For each pathway, leaky cell labeling comprised about 2% of the total cell population labeled by the other pathway. Thus, despite this potential for viral leakage, anterograde transneuronal transport was overall quite selective in labeling the expected post-synaptic granule or Purkinje cell populations.

### Functional characterization of AAV1 transsynaptic spread

To explore the association between anterograde transsynaptically labeled neurons and functional connectivity, we injected a 1:1 mixture of scAAV1-hSyn-Cre and AAV1-EF1a-DIO-ChR2-YFP into the primary auditory cortex (A1) of Ai14 mice and performed slice recording from tdTomato+ neurons and neighboring tdTomato-neurons in the inferior colliculus (IC) (Figure 2A). Expression of channelrhodopsin2 (ChR2) was restricted to neurons that were co-transduced by scAAV1-Cre and AAV1-DIO-ChR2, thus allowing us to activate the same presynaptic axons that presumably transported scAAV1-Cre to its downstream targets, which resulted in tdTomato labeling of the target IC cells (Figure 2B). Using whole-cell recording, functionally connected IC neurons were identified by their LED-evoked excitatory synaptic responses (Figure 2C), which persisted in the presence of tetrodotoxin (TTX) and 4-AP (Petreanu et al., 2009). Altogether, 100% of the tdTomato+ cells recorded (28/28 cells) exhibited LED-evoked monosynaptic excitatory responses, whereas only 46% of neighboring tdTomato-cells (13/28 cells) responded to LED (Figure 2D-E). These results revealed a statistically significant association between tdTomato+ labeling and functional connectivity when compared with non-labeled neurons randomly recorded within the same region and functionally demonstrate that AAV1 preferentially spreads to synaptically connected neurons downstream of the injection site.

**Figure 2.**
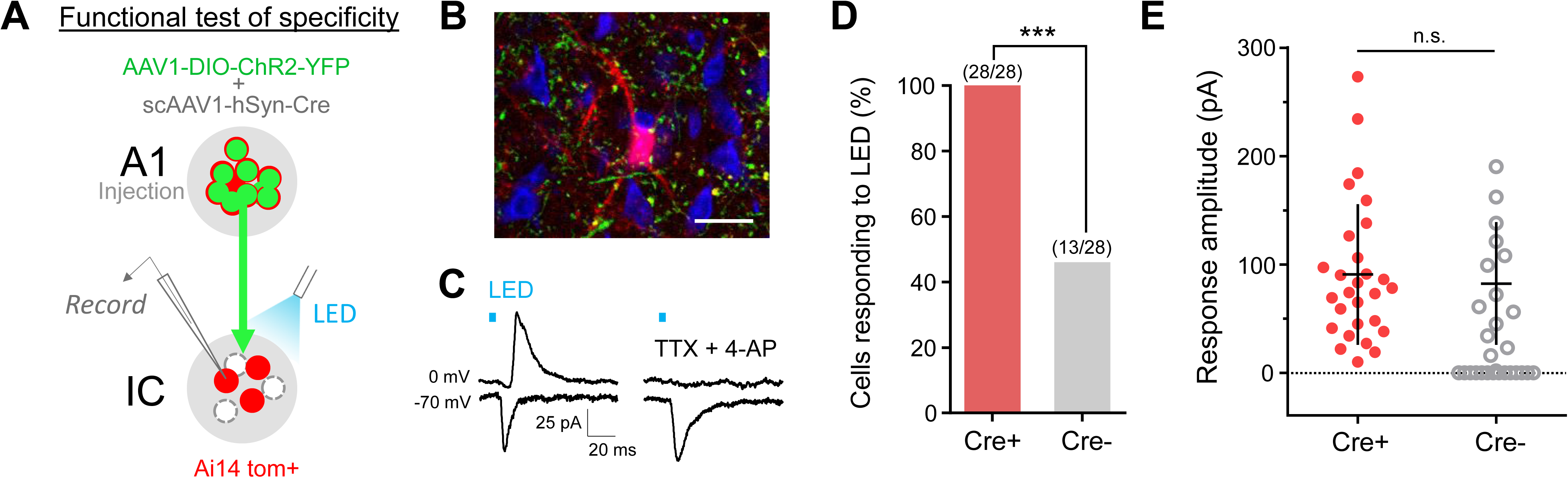
Verification of functional synaptic connectivity.

A. Strategy for slice recording from transsynaptically labeled neurons in the IC (red) following co-injection of scAAV1-hSyn-Cre and AAV1-DIO-ChR2-YFP into A1.
B. ChR2-expressing axon terminals (green) surrounding a tdTomato-labeled neuron and neighboring non-labeled neurons (blue, Nissl stain). Scale bar, 25 µm.
C. Average LED-evoked excitatory (recorded at −70 mV) and inhibitory (0 mV) currents in an example tdTomato+ IC neuron before (left) and after (right) perfusing in TTX and 4-AP. LED stimulation is marked by a blue bar.
D. Fraction of transsynaptically labeled (red) and neighboring non-labeled (gray) neurons showing monosynaptic excitatory currents in response to LED stimulation. ***p < 0.001, Chi-square test, 28 cells in each group.
E. Summary of amplitudes of average monosynaptic excitatory currents evoked by LED for all labeled (red) and non-labeled (gray) recorded neurons (neurons showing zero currents were excluded). Error bar = SD. There is no significance, unpaired t-test.

### Dependence on synaptic vesicle release

How might AAV1 be released from synaptic terminals? Previous studies have shown that, following uptake at the soma, about 14% of AAV-containing endosomes are actively trafficked down the axon in a kinesin-2 dependent manner (Castle et al., 2014a; Castle et al., 2014b). It is possible that some of these may merge with endosomal compartments that give rise to synaptic vesicles, enabling co-release of AAV particles and neurotransmitter into the synaptic cleft (Figure 3A). To examine the contribution of synaptic vesicle release in viral spread, we expressed tetanus toxin light chain (TeNT) in V1 neurons using injections of AAVDJ-CMV-TeNT-P2A-GFP (or AAV1-hSyn-GFP as control) in Ai14 mice (Figure 3A). TeNT has been shown to completely block Ca^2+^-evoked synaptic vesicle fusion and transmitter release by cleaving VAMP2 (also known as synaptobrevin-2) (Schaivo et al., 1992; Schoch et al., 2001; Yamamoto et al., 2003). Cleavage results in improper SNARE complex formation specifically for synaptic vesicles, while presumably leaving other forms of vesicular fusion intact. Following a 2-week post-injection survival time to allow for TeNT expression, a second injection of scAAV1-hSyn-Cre was targeted to the same location in V1 (Figure 3B-C, top panel), and SC was then examined 2 weeks later for the presence of tdTomato+ cell bodies (Figure 3B-C, bottom panels). Remarkably, we observed a ∼94% decrease in the number of transsynaptically labeled (i.e. tdTomato+) neurons in SC, as compared with control animals that received only GFP-expressing AAV1 injection followed by scAAV1-hSyn-Cre injection (Figure 3D). The remaining fraction of transsynaptically labeled cells observed may be the result of incomplete co-transduction of starter cells with both viruses. Lastly, it is also worth noting that the reduction in labeling of SC neurons was accompanied by noticeable enlargements in TeNT-containing axon terminals (Figure 3C, bottom panel), which may reflect a compensatory change following complete block of synaptic vesicle release (see Woods et al., 2018). Overall, these results provide additional insight into the synaptic specificity of viral spread and suggest that successful transduction of downstream neurons by AAV1 may favor a synaptic mechanism of vesicle release.

**Figure 3.**
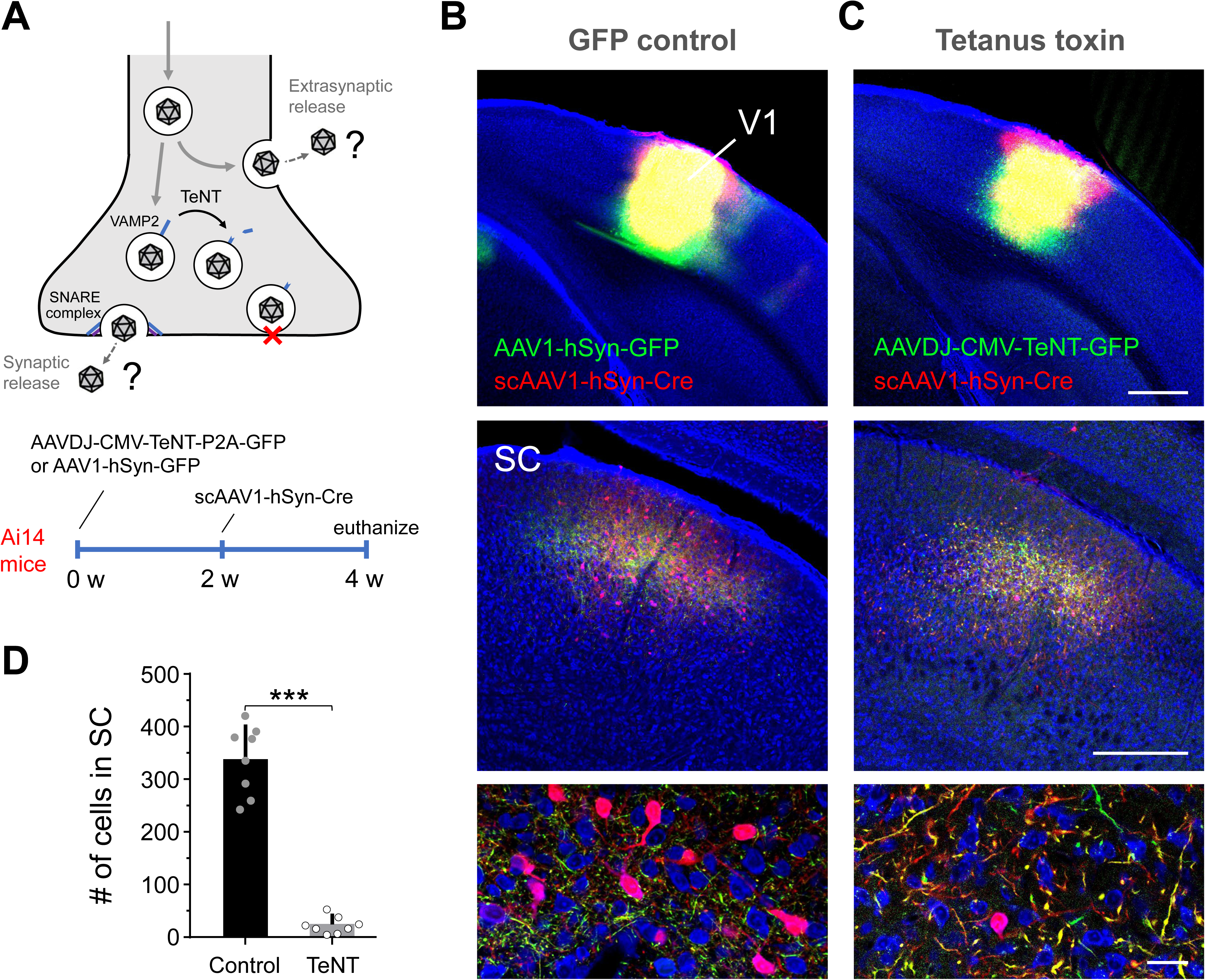
Tetanus toxin inhibition of viral spread.

A. Experimental design and timeline of virus injections. AAV trafficked to the synapse may be released through synaptic vesicles in a VAMP2-dependent manner (upper). Tetanus toxin cleaves VAMP2, preventing synaptic vesicle fusion and potential release of AAV.
B. Control experiment. AAV1-hSyn-GFP injection in V1 followed by scAAV1-hSyn-Cre injection two weeks later in Ai14 mice (top panel). Two weeks after the second injection, GFP+ axons (green) and anterograde transsynaptically labeled neurons (red cells) were observed in SC (middle panel, 10X; bottom panel, 40X). Blue, fluorescent Nissl stain.
C. AAVDJ-CMV-TeNT-P2A-GFP injection in V1 followed by scAAV1-hSyn-Cre injection two weeks later in Ai14 mice (top panel). Two weeks after the second injection, GFP+/TeNT+ axons (green) were found in SC, however very few transsynaptically labeled neurons (red cells) were observed. Scale bars (B and C): 500 µm, top panel; 250 µm, middle; 25 µm, bottom.
D. Quantification of number of anterograde transsynaptically labeled cells in SC (n = 8 mice for each group). Error bar = SD. ***p < 0.001, t-test.

### Efficiency of AAV1 spread through inhibitory synapses

We previously demonstrated that AAV1-Cre can spread from excitatory projection neurons to both excitatory and inhibitory postsynaptic cell-types (Zingg et al., 2017). However, whether or not this transport occurs efficiently across other types of synaptic connections in the brain (e.g. through inhibitory or neuromodulatory pathways) remains to be determined. To test this, we first examined the efficiency of anterograde transsynaptic spread from inhibitory neurons to downstream excitatory or inhibitory cell populations in Ai14 mice. To avoid possible retrograde labeling from AAV1 injections (Rothermel et al., 2013; Aschauer et al., 2013; Zingg et al., 2017), only unidirectionally connected regions downstream of each injection site were examined. To characterize the efficiency of spread from inhibitory-to-inhibitory cells, we injected scAAV1-hSyn-Cre into the striatum (Str), which contains GABAergic medium spiny neurons (MSNs) that project to inhibitory cells within the substantia nigra, pars reticulata (SNr) (Figure 4A, left two panels). Following a 2-week post-injection survival time, numerous tdTomato+ cells were found intermingled with dense axon terminals in SNr (Figure 4A, middle panels), suggesting the potential for AAV1 to spread through inhibitory synapses to downstream inhibitory neurons. We also observed tdTomato+ cells in the overlying substantia nigra, pars compacta (SNc, presumed dopaminergic neurons) (Figure 4A, third from left panel), however these were excluded from analysis as they provide strong projections back to the Str and may therefore have been retrogradely labeled by AAV1-Cre. This is supported by the observation that injections of GFP-expressing G-deleted rabies virus into the same region of the Str robustly back-label neurons in SNc, but not in SNr (Figure 4A, right panel). To provide an estimate of the anterograde transsynaptic labeling efficiency from Str→SNr, we quantified the number of tdTomato+ neurons relative to the total number of Nissl+ neurons within the boundaries of the axon terminal field in SNr (see Materals and Methods). We found on average ∼41% of the cells within this region were tdTomato+ (Figure 4F, grey), suggesting comparable efficiency to previously reported excitatory projection neuron pathways (Zingg et al., 2017). In addition, using the same approach, we also tested AAV1 spread from inhibitory neurons in the SNr to presumed excitatory neurons in the ventromedial nucleus of the thalamus (VM), another known unidirectional pathway (Figure 4B). Again, we found relatively efficient anterograde transsynaptic labeling in VM (∼36% tom+/total Nissl+ cells; Figure 4F, light gray), providing further evidence that AAV1 may be used to transsynaptically tag diverse cell populations downstream of different classes of inhibitory projection neurons.

**Figure 4.**
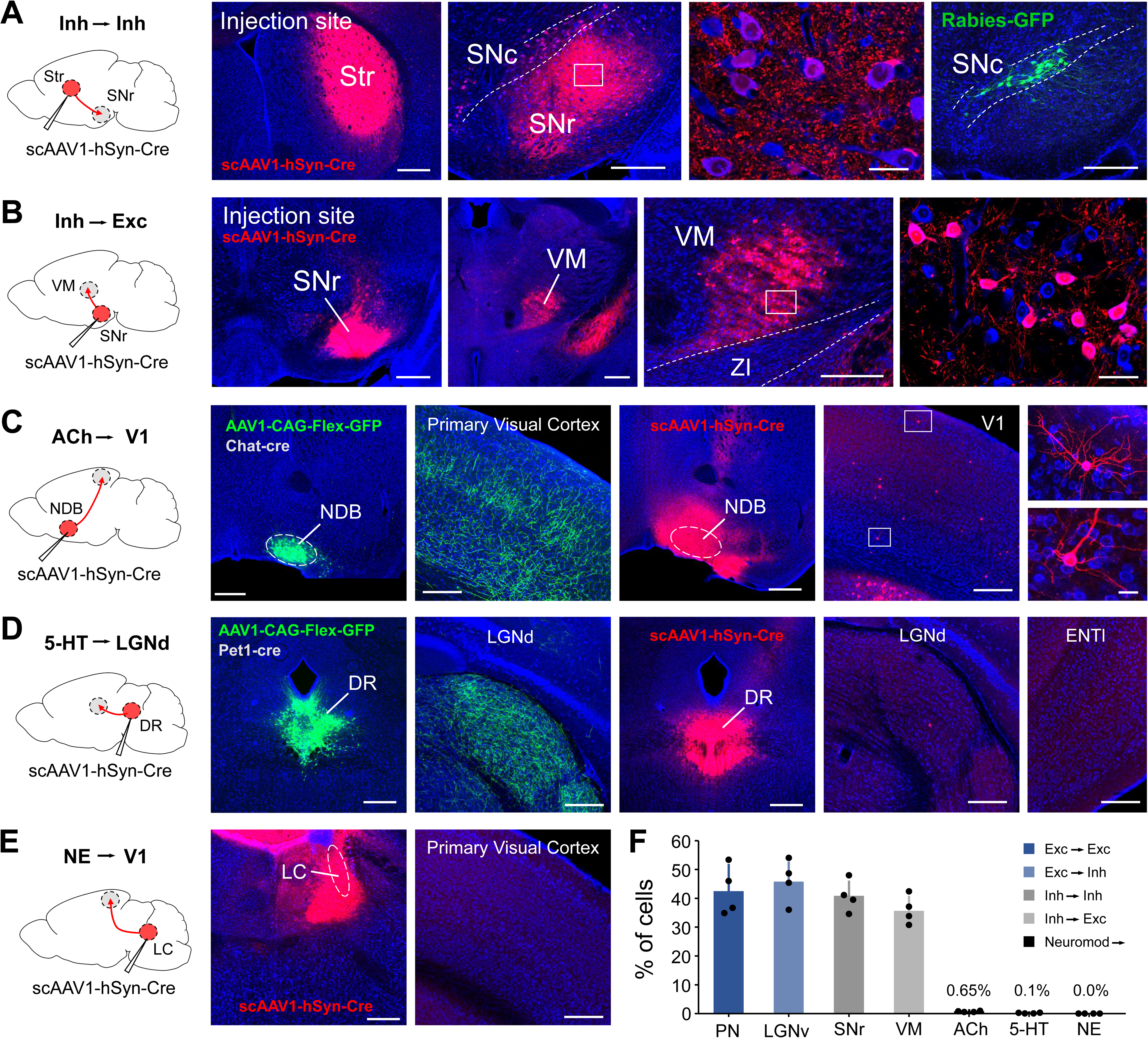
Efficiency of viral spread across different types of synapses.

A. Anterograde transsynaptic spread of scAAV1-hSyn-Cre in Ai14 mice from inhibitory projection neurons in striatum (left two panels, red, scale bar, 500 µm) to inhibitory neurons in the SNr (middle two panels, scale bars, 250 µm, left; 25 µm, right). Retrograde labeling by Rabies-GFP injection in striatum was restricted primarily to SNc (right panel, green, scale bar, 250 µm).
B. Anterograde transsynaptic spread of scAAV1-hSyn-Cre from inhibitory projection neurons in SNr (left two panels, red, scale bar, 500 µm) to presumed excitatory neurons in VM (right three panels, scale bars, 500 µm, left; 250 µm, middle; 25 µm, right).
C. Anterograde transsynaptic spread through cholinergic neurons (left panel). Cholinergic cells in the NDB project unidirectionally to the primary visual cortex (left second and third panels, green, scale bars, 500 µm, left; 250 µm, right). Injection of scAAV1-hSyn-Cre into the NDB sparsely labeled neurons in V1 across different cortical layers (right three panels, red, scale bars, 500 µm, left; 250 µm, middle; 25 µm, right).
D. Anterograde transsynaptic spread through serotonergic neurons (left panel). Serotonergic neurons in DR project unidirectionally to LGNd (left second and third panels, green, scale bars, 500 µm, left; 250 µm, right) and ENTl (data not shown). Injection of scAAV1-hSyn-Cre into the DR labeled little or no cells in LGNd or ENTl (right three panels, red, scale bars, 500 µm, left; 250 µm, middle; 250 µm, right).
E. Anterograde transsynaptic spread through noradrenergic neurons. Injection of scAAV1-hSyn-Cre into the LC (left two panels, red, scale bar, 500 µm). No labeling of cell bodies was observed in downstream regions unidirectionally connected to LC, such as V1 (right panel, scale bar, 250 µm).
F. Efficiency of transsynaptic spread across different types of synapses (# of tdTomato+ cells / # of Nissl+ cells) in each target region shown in (A-E) (n = 4 mice each). Data are compared with previous results for excitatory projections from V1 to downstream excitatory (PN) and inhibitory (LGNv) cell-types (blue bars). Error bar = SD.

### Efficiency of AAV1 spread across neuromodulatory synapses

To determine whether neuromodulatory cell-types support anterograde transsynaptic spread of AAV1, we performed similar injections in brain regions that contain cholinergic (ACh), serotonergic (5-HT), or noradrenergic (NE) cell populations and examined targets that were exclusively downstream for tdTomato+ cell bodies (Figure 4C-E). Specifically, to reveal transsynaptic spread through cholinergic neurons, we injected the diagonal band nucleus (NDB) in the anterior basal forebrain, which projects strongly to V1 (Figure 4C, second and third from left panels), but does not receive input back from V1. Following injections of scAAV1-hSyn-Cre into the NDB, we observed sparse tdTomato+ neurons scattered throughout all layers of V1, including layer 1 (Figure 4C, right three panels). This suggests AAV1 may have the capacity to spread post-synaptically through cholinergic neurons, albeit much less efficiently than expected given the dense termination pattern in this target structure (Figure 4C, third from left panel). Indeed, quantification of the number of tdTomato+ cells relative to Nissl+ cells within a given 500 × 500 µm region of V1 revealed only ∼0.65% labeling efficiency (Figure 4F). Similarly, we also observed inefficient spread of AAV1 through serotonergic and noradrenergic cell populations. Serotonergic projections from the dorsal raphe (DR) to the dorsolateral geniculate nucleus (LGNd) or lateral entorhinal cortex (ENTl) produced almost no transsynaptic labeling (Figure 4D, right three panels), despite strong axonal innervation of these structures (Figure 4D, second and third from left panels). Furthermore, transsynaptic labeling was completely absent in V1 following transduction of noradrenergic neurons in the locus coeruleus (LC) (Figure 4E), which diffusely project to most cortical areas, including V1 (Polack et al., 2013; Schwarz et al., 2015). Thus, while AAV1 appears to be capable of efficient spread through classically defined glutamatergic and GABAergic synaptic pathways, its application in various neuromodulatory systems may be limited.

### Transsynaptic categorization of input-defined spinal cord neurons

We next asked whether AAV1 transsynaptic tagging could be applied in long-range projection pathways from the brain to the spinal cord. To test this, we first identified several brain regions that contained spinal-projecting cell populations by injecting retrogradely transported AAV (AAVretro; Tervo et al., 2016) expressing Cre or GFP in the cervical and lumbar spinal cord, respectively, of Ai14 mice. Based on the retrograde labeling result (Figure 5A), we selected two cortical regions (primary motor cortex, MOp; and somatosensory cortex, SSp) and three subcortical regions (lateral hypothalamic area, LHA; red nucleus, RN; and superior vestibular nucleus, SUV) for injections of scAAV1-hSyn-Cre (Figure 5B). Following a 2-week post-injection survival time, robust tdTomato+ cell body labeling was observed in the spinal cord for each descending pathway (Figure 5C), suggesting a capacity for AAV1-Cre to efficiently transduce neurons over long distances. Interestingly, the distribution of this postsynaptic labeling appeared to be regional and layer-specific for each pathway, suggesting that different descending projections to the spinal cord recruit unique sub-populations of spinal neurons that may underlie their distinct functional roles. For example, MOp labeled neurons were in lamina IV and V of the spinal cord, respectively (Figure 5C, first column from left), while output from SSp was restricted mostly to the medial portion of lamina IV (Figure 5C, second column from left). These results further highlight the specificity of anterograde transsynaptic AAV1 spread and reveal its potential use in dissecting the downstream circuit components of different brain-spinal projection pathways.

**Figure 5.**
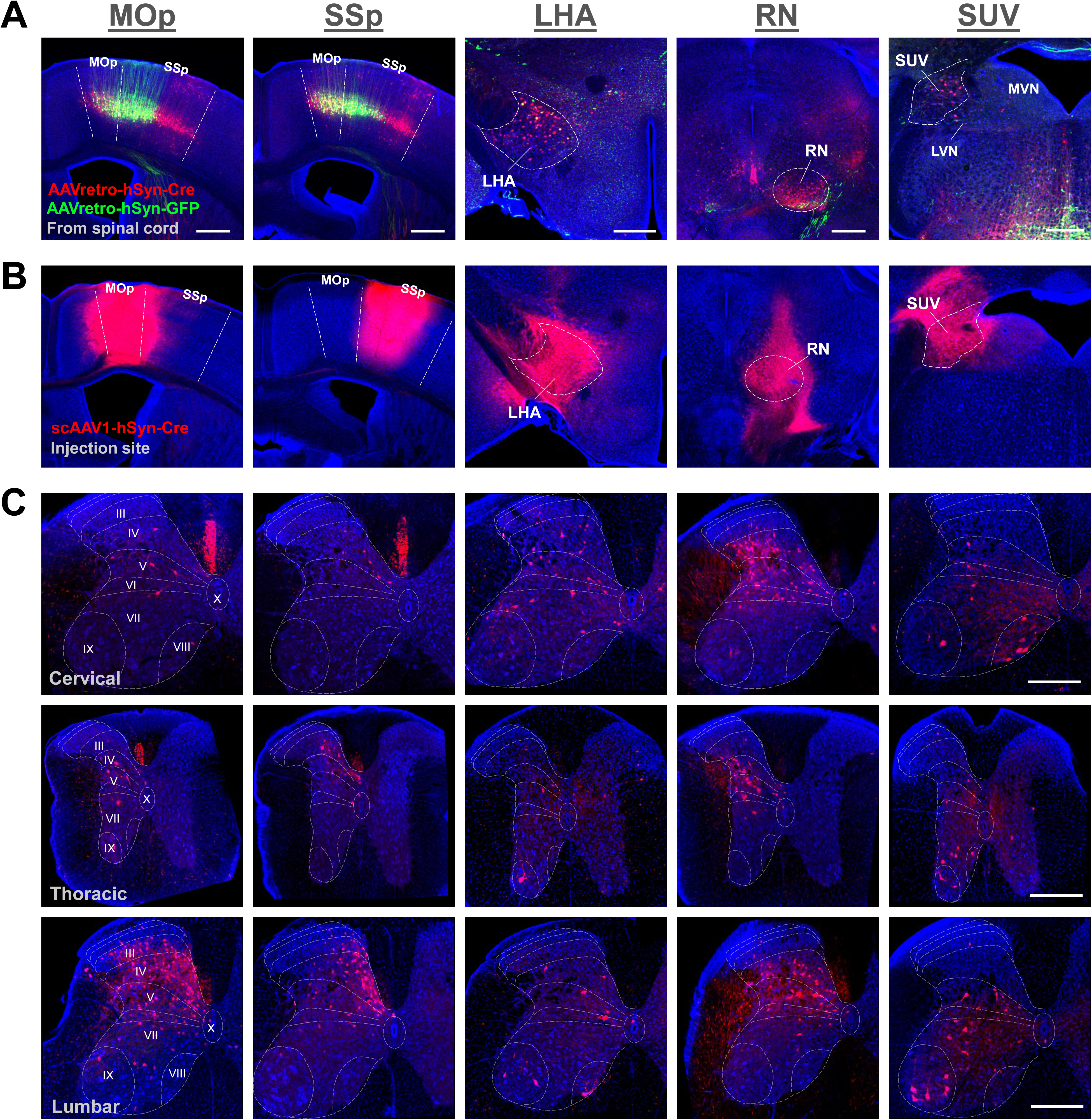
Application in descending pathways to the spinal cord.

A. Retrograde labeling of spinal-projecting cortical and subcortical neuronal populations following injections of AAVretro-hSyn-Cre (red) and AAVretro-hSyn-GFP (green) into the left side of the cervical and lumbar spinal cord, respectively, in Ai14 mice. Scale bars (A and B), 500 µm.
B. Corresponding injections of scAAV1-hSyn-Cre (red, 100 nL injection volume) into different spinal-projecting brain regions (marked on top of panel A) in Ai14 mice.
C. Different patterns of transsynaptic labeling at cervical, thoracic, and lumbar segments of the spinal cord for each injection following a 2 week post-injection survival time. Each column corresponds to the injection site shown above in (B). Scale bars, 250 µm. Abbreviations: LHA, lateral hypothalamic area; LVN, lateral vestibular nucleus; MOp, primary motor cortex, lower limb;; MVN, medial vestibular nucleus; RN, red nucleus; SSp, primary somatosensory cortex; SUV, superior vestibular nucleus.

### Dual anterograde transsynaptic tagging of topographically defined cell populations

As corticofugal output represents one of the largest and most straight-forward systems for applying anterograde transsynaptic tagging, we aimed to explore in greater detail its potential for accessing input-defined cell populations throughout the entire brain. In particular, given the highly topographical nature of corticofugal output, we asked whether AAV1-Cre and AAV1-Flp could be used together to simultaneously reveal the topographic distribution of cells innervated by two functionally distinct cortical regions. To test this, we injected AAV1-hSyn-Flp in MOp-ul and scAAV1-hSyn-Cre into MOp-ll in a Flp- and Cre-reporter mouse (Frt-GFP × Ai14-tdTomato) (Figure 6A-B). After 2 weeks, we then examined all subcortical targets for the presence of GFP+/Flp+ or tdTomato+/Cre+ cell bodies corresponding to output from either MOp-ul or MOp-ll, respectively. The thalamus was excluded from analysis due to its reciprocal connectivity with each injection site, however topography was still evident in this structure (Figure 6C, top row, third panel from left). Remarkably, several brain regions, including the striatum (Str), zona incerta (ZI), medial accessory oculomotor nucleus (MA3), anterior pretectal nucleus (APN), RN, PN, and SC, contained discrete, non-overlapping populations of GFP+ or tdTomato+ cells that subdivided each structure based on its input from upper- or lower-limb-related motor cortex (Figure 6C). In addition, we also observed structures such as the bed nucleus of the anterior commissure (BAC), caudal periaqueductal gray (PAG), pontine reticular nucleus (PRN), and cuneate (CU) and gracile (GR) nuclei, that were preferentially labeled by one pathway, but not the other (Figure 6C).

**Figure 6.**
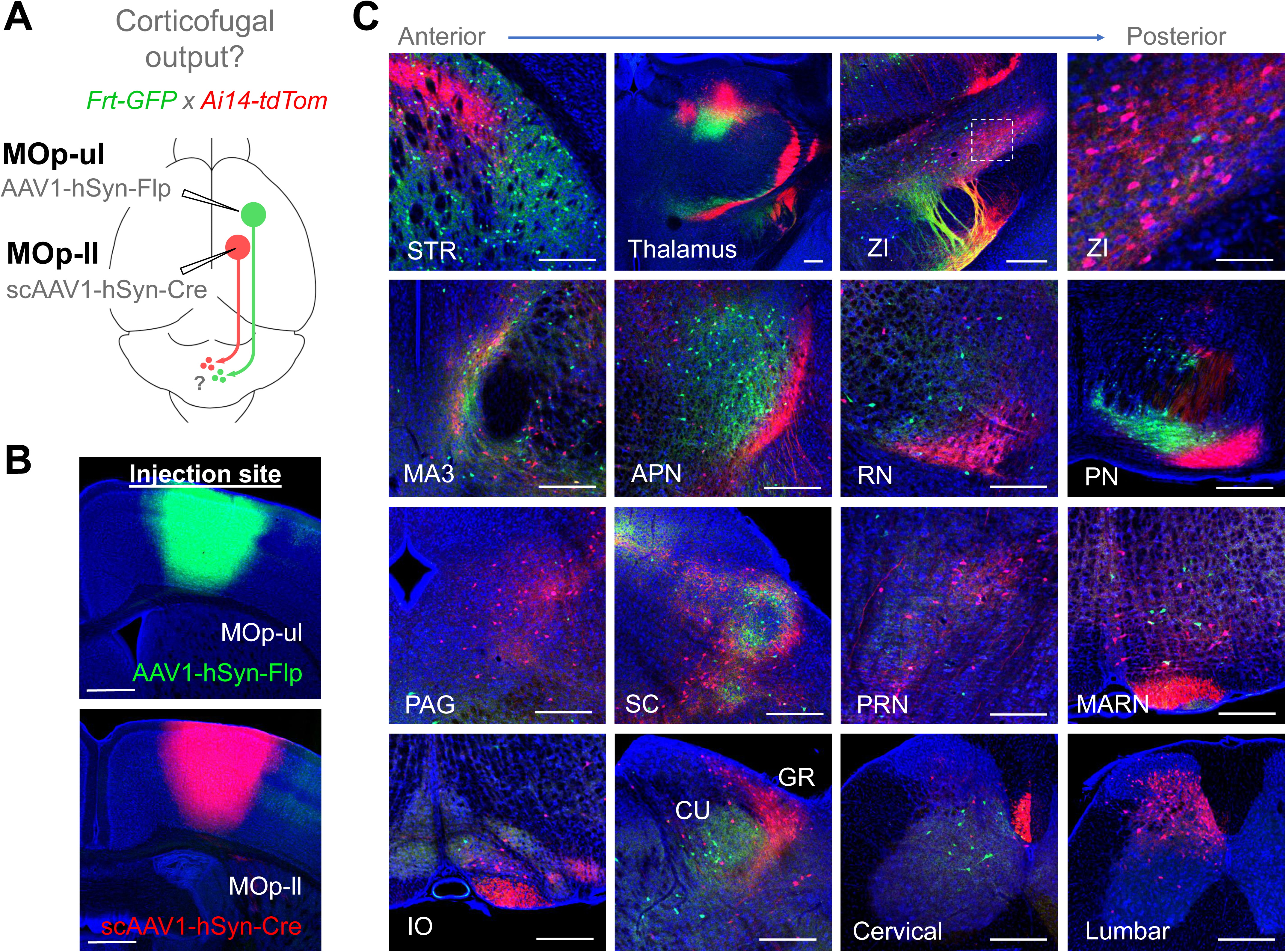
Accessing topographically precise, input-defined cell populations through corticofugal pathways.

A. Strategy for labeling cell populations that receive input from upper limb- (green) and lower limb- (red) related primary motor cortex in Ai14 × Frt-GFP (Cre and Flp reporter) mice.
B. Injection sites for AAV1-hSyn-Flp in MOp-ul (top panel, green) and scAAV1-hSyn-Cre in MOp-ll (bottom panel, red) in an Ai14 × Frt-GFP mouse. Scale, 500 µm.
C. Anterograde transsynaptic labeling of cells that receive input from upper limb- (green) or lower limb- (red) related primary motor cortex. Many closely apposed, non-overlapping cell populations were observed in mid- and hindbrain structures, including the PN (second row, right panel). None of the structures shown project back to motor cortex, with the exception of the thalamus (top row, third panel from left), which may contain both retrograde and anterograde transsynaptic labeling of cell bodies. Scale bars, 250 µm. Abbreviations: APN, anterior pretectal nucleus; BAC, bed nucleus of the anterior commissure; CU, cuneate nucleus; GR, gracile nucleus; IO, inferior olive; MA3, medial accessory oculomotor nucleus; MARN, magnocellular reticular nucleus; PAG, periaqueductal gray; PN, pontine nucleus; PRN, pontine reticular nucleus; RN, red nucleus; SC, superior colliculus; STR, Striatum; ZI, zona incerta.

Given this potential to subdivide certain structures based on their topographic input (Figure 7A), we next asked whether PN neurons defined by their input from MOp-ul or MOp-ll might in turn project to upper- or lower-limb-related portions of the cerebellum, thus bridging two somatotopically organized systems and providing some cross-validation for the specificity of these transsynaptically tagged subpopulations. To test this, we injected scAAV1-hSyn-Cre into MOp-ul or MOp-ll and AAV1-CAG-Flex-GFP into the PN to Cre-dependently express GFP in each subpopulation of input-defined PN neurons (Figure 7B,E). We then examined the cerebellum for GFP+ axon terminals in each case. Interestingly, we found that MOp-ul-recipient PN neurons projected primarily to the contralateral paramedian lobule (PRM), which has been shown to respond to forelimb stimulation in micromapping studies (Shambes et al., 1978; Odeh et al., 2005) and has been shown to receive di-synaptic input specifically from the forelimb motor cortex via the pontine nucleus using multi-synaptic rabies virus tracing (Suzuki et al., 2012) (Figure 7C-D). On the other hand, MOp-ll-recipient PN neurons projected strongly to the contralateral copula pyramidis (COP) lobule of the cerebellum, but avoided the PRM (Figure 7F-H), as expected based on previous functional and anatomical mapping studies (Atkins & Apps, 1997; Voogd et al., 2003; Odeh et al., 2005; Suzuki et al., 2012). Together, these results provide evidence that AAV1-transsynaptic tagging may be broadly applied in various corticofugal pathways to experimentally access, map, and manipulate specific subpopulations of neurons defined by their topographic cortical input.

**Figure 7.**
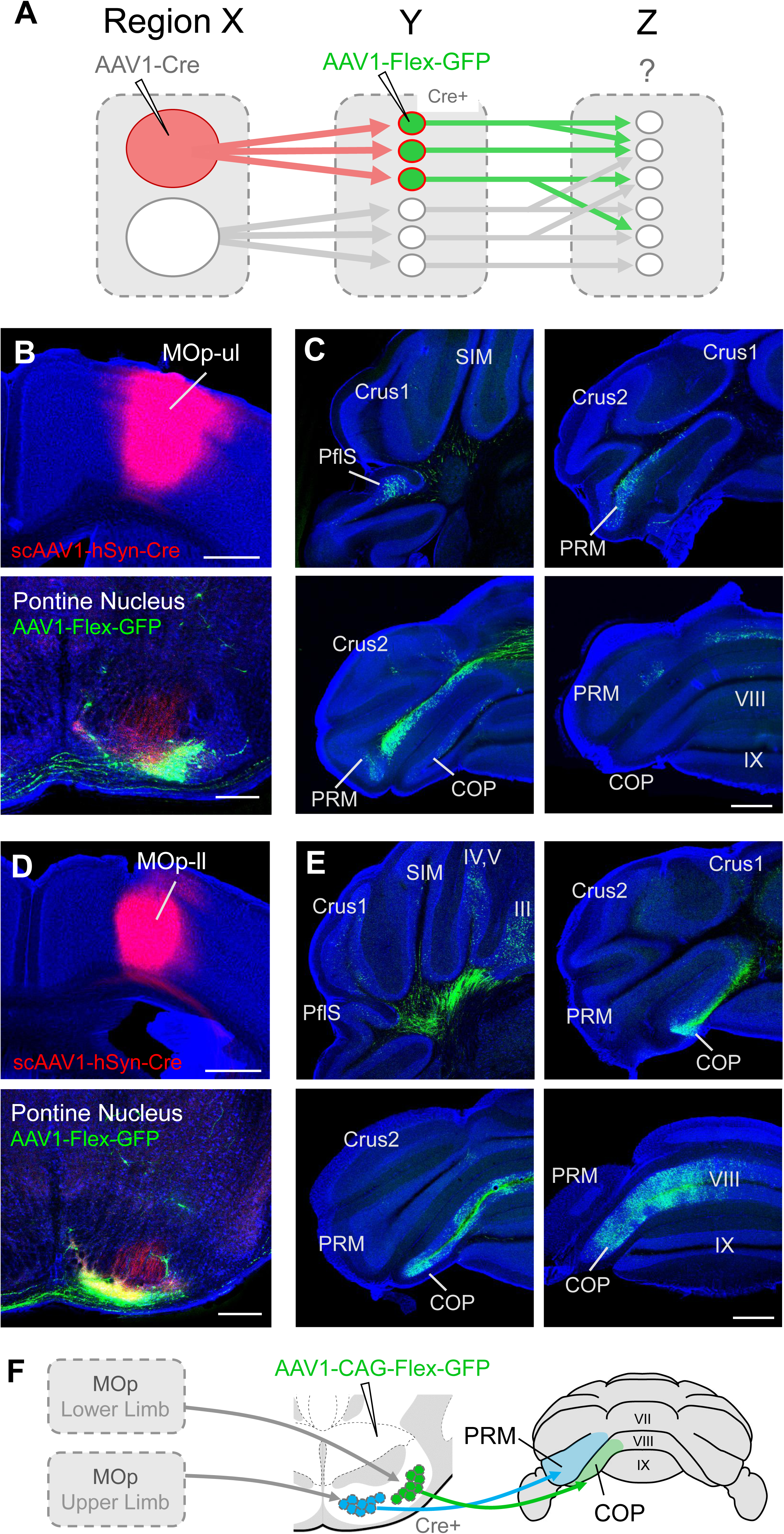
Forward mapping of topographically organized brain regions.

A. Schematic diagram depicting the parcellation of brain regions (in Y) based on their topographical input (from X) using an anterograde transsynaptic approach. Secondary injections of AAV1-Flex-GFP enable further mapping of axonal outputs (to Z) for any given input-defined subregion. A retrograde approach may not provide access to the same population due to collateralization of divergent outputs.
B. Example of topographic mapping of PN neurons, which receive spatially distinct input from MOp-ul and MOp-ll (see Figure 6C, second row, right panel). Cre+/GFP+ neurons are shown in PN following injection of scAAV1-hSyn-Cre in MOp-ul and AAV1-hSyn-Flex-GFP in PN. Scale bar, 250 µm.
C. Axonal projections to the cerebellum from PN neurons defined by their input from MOp-ul. Axons primarily target the copula pyramidis (COP, top right and bottom panels), but also collateralize more sparsely to lobules III and IV,V (top left panel). Scale bar, 500 µm.
D. 10X magnification of mossy fibers shown in the boxed region in (C). Scale bar, 250 µm.
E. Cre+/GFP+ neurons in PN following injection of scAAV1-hSyn-Cre in MOp-ll and AAV1-hSyn-Flex-GFP in PN. Scale bar, 250 µm.
F. Axonal targeting in cerebellar cortex. Output was primarily restricted to contralateral parafloccular sulcus (PflS, top left panel) and the ventral paramedian lobule (PRM, bottom left panel). Scale bars, 500 µm.
G. 10X magnification of axons shown in the boxed region in (F). PRM targeting appeared as segregated bands. Scale bar, 250 µm.
H. Schematic summary of axonal projections to cerebellum from MOp-ul and MOp-ll-recipient PN neurons (green and blue, respectively). Posterior view of the cerebellum is shown.

### Application in sparse labeling approaches for single neuron reconstruction

To better characterize populations of neurons that receive input from a given pathway, it may be useful to recover their individual morphological and axonal targeting features. Given the normally dense arrangement of these processes, this is greatly facilitated by using a method to sparsely label only a small number of cells at a time. To achieve sparse labeling in a given Cre-expressing cell population, we reasoned that co-injections of a low titer Cre-dependent Flp-expressing virus (AAV1-DIO-Flp) and a high titer Flp-dependent YFP-expressing virus (AAVDJ-fDIO-YFP) could be used to obtain robust levels of YFP expression in only a few cells (Figure 8A), similar to a recent study (Lin et al., 2018). To establish a relationship between titer and the resulting number of labeled cells, AAV1-DIO-Flp was diluted to a final concentration of either 7.5 × 10^8^, 10^9^, or 10^10^ GC/mL and co-injected with AAVDJ-fDIO-YFP (1.2 × 10^13^ GC/mL, 50 nL total volume) in V1 of Ai14 × PV-Cre mice, which express Cre in parvalbumin (PV)+ cortical interneurons (Figure 8B). At this injection volume, we found that titers around 7.5 × 10^9^ GC/mL consistently labeled ∼4 PV+ cells per animal, enabling recovery of cell morphology (Figure 8C), while titers around 7.5 × 10^8^ GC/mL labeled only 1 cell in one out of four animals (Figure 8B,D). Therefore, working around a concentration of 10^9^ GC/mL and adjusting for the relative difference in starter population density, it may be possible to apply this approach to any given group Cre-expressing neurons, including those labeled using anterograde transsynaptic spread of AAV1-Cre.

**Figure 8.**
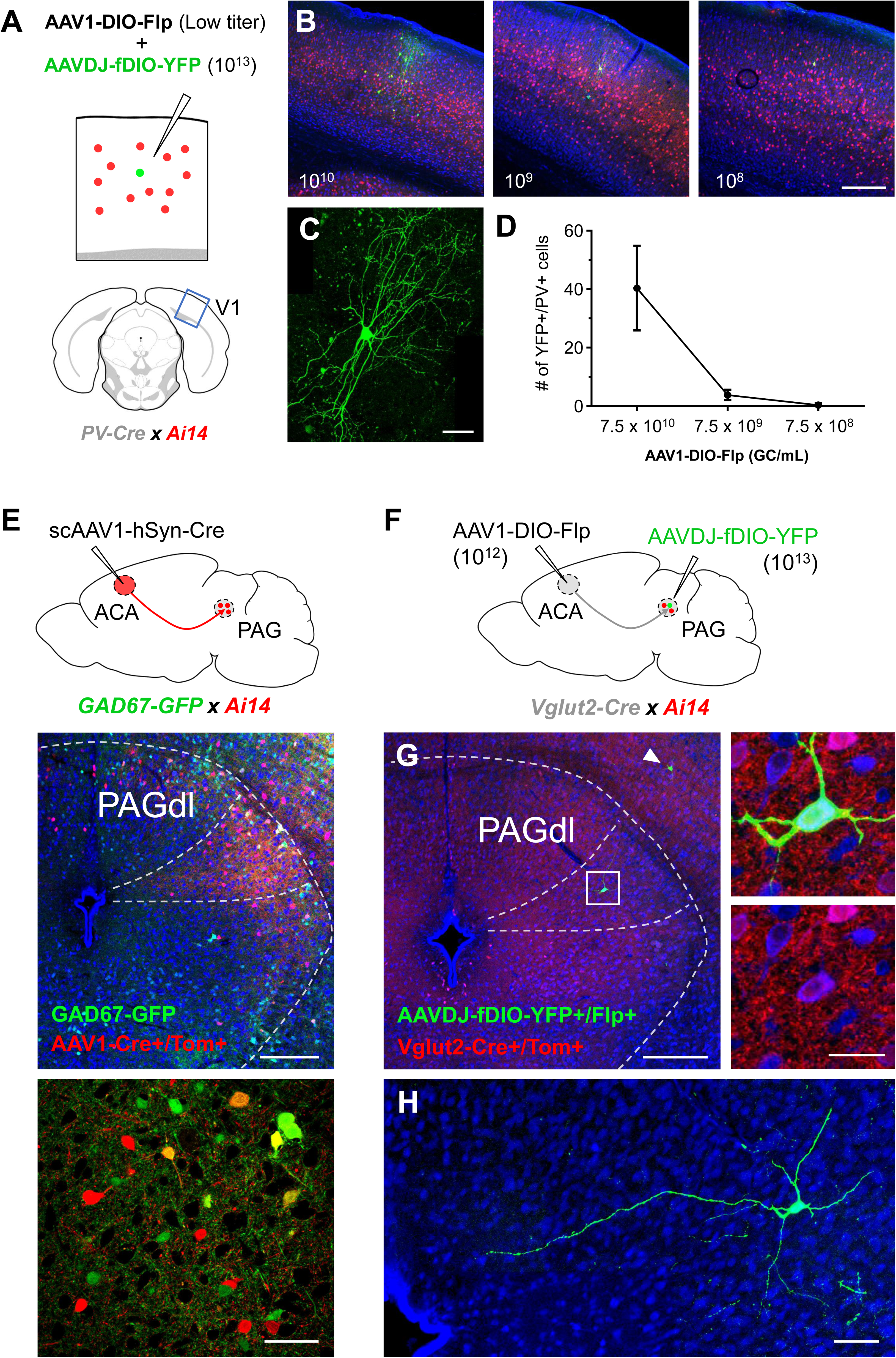
Application with sparse labeling approaches for reconstructing single neuron morphology.

A. For a given Cre+ cell population (red), sparse labeling (green) may be achieved by co-injecting AAV1-DIO-Flp at increasingly lower titers along with high titer AAVDJ-fDIO-YFP. To establish a titering curve, PV neurons in V1 were targeted with co-injections of AAVDJ-fDIO-YFP (final titer: 1.2 × 10^13^ GC/mL) and AAV1-DIO-Flp (final titers: 7.5 × 10^10^, 10^9^, or 10^8^ GC/mL) in PV-Cre × Ai14 mice.
B. Examples of YFP cell labeling (green) achieved at each titer step. Red, PV-Cre+/tdTomato+ cells. Blue, fluorescent Nissl. Scale bar, 250 µm.
C. 40X magnification of a YFP+ PV neuron labeled in (B, middle panel). Scale bar, 25 µm.
D. Quantification of the number of YFP+/PV+ cells labeled at each titer step (n = 4 mice each). Error bar, SD.
E. Injection of scAAV1-hSyn-Cre in the ACA of GAD67-GFP × Ai14 mice transsynaptically labels neurons in PAG. Labeled cell density is greatest in the dorsolateral PAG (PAGdl, middle panel), and both inhibitory (GFP+/tdTomato+) and presumed excitatory (GFP-/tdTomato+) cell-types are labeled (bottom panel, 40X magnification). Scale bars, 250 µm, middle panel; 50 µm, bottom panel.
F. Strategy for sparse labeling of input- and genetically-defined cell populations. AAV1-DIO-Flp titer for injection in ACA is reduced to 1.5 × 10^12^ GC/mL to achieve sparse anterograde transsynaptic labeling in PAGdl. Individual Vglut2-Cre+/Flp+ neurons may then be targeted with an injection of high titer AAVDJ-fDIO-YFP to specifically label glutamatergic neurons in PAGdl that receive input from ACA.
G. Example of a single Vglut2-Cre+/Flp+ neuron labeled in PAGdl (green). An additional neuron was found in the superior colliculus (arrowhead). Right panels, 40X magnification of YFP+ (green, upper panel) and Vglut2-Cre+/Tom+ (red, lower panel) neuron in PAGdl. Blue, fluorescent Nissl. Scale bars, 250 µm, left panel; 25 µm, right panels.
H. 40X magnification of local dendrites and axonal projection of the YFP+ PAGdl neuron shown in (G). Scale bar, 50 µm.

Transsynaptic tagging may be applied in different Cre-expressing transgenic mice to access both input- and genetically-defined cell-types in a given downstream circuit (Zingg et al., 2017). We therefore explored an additional means to achieve sparse labeling in cell populations that meet these criteria. As a test system, we examined the projection from the anterior cingulate cortex (ACA) to the dorsolateral periaqueductal gray (PAGdl) (Figure 8E). Following injections of scAAV1-hSyn-Cre in the ACA of Ai14 × GAD67-GFP mice, a mixture of transsynaptically labeled GABAergic (GFP+/tdTomato+) and presumed glutamatergic (GFP-/tdTomato+) neurons were observed primarily within the PAGdl (Figure 8E, bottom panels), suggesting descending projections from ACA may innervate both cell-types. To select only glutamatergic neurons for analysis, similar injections may be performed in ACA of Vglut2-Cre mice using AAV1-DIO-Flp and a secondary injection of AAVDJ-fDIO-YFP within the PAGdl to label input-defined glutamatergic Cre+/Flp+ cells with YFP. With high titer AAV1-DIO-Flp (e.g. >10^13^ GC/mL) and AAVDJ-fDIO-YFP, these injections are expected to label a large number of cells in PAGdl. However, AAV1-DIO-Flp may be diluted to achieve less transsynaptic labeling, as we previously demonstrated that efficiency of spread is titer-dependent (Zingg et al., 2017). In particular, previous results revealed that reducing viral titer from ∼10^13^ GC/mL to ∼10^12^ GC/mL reduces transsynaptic labeling efficiency by about 85-90%. As our test injection of scAAV1-hSyn-Cre at 10^13^ titer yielded an average of 35 GFP-/tdTomato+ cells (presumed glutamatergic neurons) per 300 µm^3^ sample space of PAGdl, we expected that similar injections of AAV1-DIO-Flp at 10^12^ titer would tag ∼4 Vglut2-Cre+/Flp+ within the same sample region (see Materials and Methods for details). This sparse population could then be targeted for robust YFP expression following local injection of high titer AAVDJ-fDIO-YFP (Figure 8F). Following this protocol, we were able to label a single input-defined glutamatergic neuron in PAGdl (Figure 8G-H). Combined with methods for tissue clearing and whole-brain imaging, it may be possible to systematically reconstruct both local morphology and all long-range axonal projections for any given sparsely labeled group of input- and genetically-defined cells.

## Discussion

In this study, we provide a systematic characterization of the synaptic specificity of AAV1 transneuronal spread and examine its transport efficiency throughout a diverse set of neural pathways. Overall, we find the virus to be highly selective in its transduction of postsynaptic neurons and broadly applicable in all pathways tested, with the exception of several neuromodulatory cell-types (i.e. serotonergic, cholinergic, and noradrenergic neurons). In addition, we reaffirm the observation that anterograde transsynaptic spread is unique to AAV1 (and to some extent AAV9), following an expanded comparison with additional AAV serotypes and other common neurotropic viruses (see Supplementary Figure 1). As expected, transsynaptic spread is not specific to Cre-expressing AAV1, *per se*, as shown by Flp-expressing virus (AAV1-hSyn-Flp) in a Flp-reporter mouse (Frt-GFP; Sousa et al., 2009). Thus, AAV1-mediated transsynaptic tagging may be expanded to include a variety of recombinase systems to facilitate intersectional approaches for accessing cell-types based on multiple criteria.

### Synaptic specificity of viral spread

We intended to address the synaptic specificity of AAV1 spread from anatomical, functional and molecular aspects. To anatomically test the synaptic specificity of AAV1 spread, we examined its capacity to exclusively label cell-types known to be innervated by either pontine or inferior olive afferents to the cerebellum (Palay & Chan-Palay, 1974; Kanichay & Silver, 2008; Mathews et al., 2012). AAV1 transport through each of these pathways demonstrated a high degree of synaptic specificity, labeling either the expected granule cell population, but not Purkinje cells (PN pathway), or Purkinje cells, but not granule cells (IO pathway).

To functionally examine the transsynaptic spread, we recorded from both transsynaptically labeled and neighboring non-labeled neurons to establish a statistical association between labeling and pre-synaptic connectivity, and between non-labeled neurons and a lack of pre-synaptic connectivity. We found that all of the labeled neurons we recorded from were mono-synaptically connected to their presynaptic partners, while only ∼46% of neighboring non-labeled cells exhibited pre-synaptic input. This revealed a highly significant association between post-synaptic AAV spread and pre-synaptic connectivity, suggesting AAV may favor a synaptic mechanism of transport.

Lastly, the dependence of transsynaptic spread on synaptic vesicle release, as demonstrated by the great impairment of the spread by TeNT expression in starting cells, suggests a predominately presynaptic mechanism of viral release. Given a potential synaptic vesicle mechanism for viral release, it would be interesting to know if neural activity is required for viral spread or if it is more efficient in cells that exhibit higher firing rates or are tonically active.

Together, our results collectively provide deeper insight into the synaptic nature of AAV1 transneuronal spread and suggest that transport through transsynaptic mechanisms contributes to the great majority of the observed labeling in each of the pathways examined here.

### Efficiency of spread through diverse brain pathways

Previously, we characterized the transsynaptic spread of AAV1 through glutamatergic projection pathways (e.g. corticofugal, retino-collicular, colliculo-thalamic). Here we provide a more systematic characterization of AAV1 spread through pathways utilizing different types of synapses (e.g. inhibitory or neuromodulatory) or projecting over long distances (e.g. cortex to spinal cord). Our results indicate that AAV1 is capable of transducing both excitatory and inhibitory cell-types downstream of GABAergic projection pathways (Str → SNr, and SNr → VM) with efficiencies comparable to previously reported glutamatergic pathways. In addition, AAV1 transport through long-range spinal-projecting populations yielded robust and regionally-specific labeling patterns in the spinal cord for each pathway tested, suggesting that transport efficiency is not compromised by axon length. On the other hand, injections in three different neuromodulatory cell populations (cholinergic, serotonergic, and noradrenergic) failed to yield efficient transsynaptic labeling in their respective downstream targets. Overall, our results suggest that AAV1 is capable of efficient transsynaptic transport through a wide variety of excitatory and inhibitory projection pathways.

### Application and future directions

In its current form, anterograde transsynaptic spread of any recombinase-expressing AAV1 may be used to gain experimental access to downstream cell populations that might otherwise be difficult to target due to their small size, laminar organization, and/or lack of identified genetic markers and corresponding transgenic mice. Given the non-toxic nature of AAV, these populations may then be targeted for long-term expression of genetically encoded tools for mapping their connectivity and probing their functional role. In addition, our results highlight the value of applying an anterograde transsynaptic approach, rather than a genetic or retrograde viral approach, in subdividing topographically organized brain regions based on their input. These regions, such as the striatum or pontine nucleus, contain relatively homogeneous distributions of cell-types, however discrete populations within these structures may process different types of information based on the source of their cortical input (e.g. from upper-limb or lower-limb related motor cortex). In turn, each population may have topographically unique output, as demonstrated in the cerebellar targeting of two input-defined subregions of PN (Figure 7H). Accessing these same populations using a retrograde viral approach, however, may not be feasible given the potential for divergent collateralization of their output (Figure 7A), and genetically specifying such input-defined populations may not be possible given the homogeneity of cell-types within the structure. Lastly, we demonstrate that AAV1 transsynaptic tagging may be used to catalog cell-types based on their input, morphology, and gene expression by incorporating the use of transgenic mice and techniques for achieving sparse labeling of transsynaptically tagged neurons (Gouwens et al., 2019; Kim et al., 2017; Li et al., 2019; Winnubst et al., 2019).

Given that AAV1 is also capable of retrograde transport (Aschauer et al., 2013; Rothermel et al., 2013; Masamizu et al., 2014; Tervo et al., 2016; Zingg et al., 2017), its application as an anterograde transsynaptic tool must be limited to unidirectional pathways. In addition, there is currently no way to initiate AAV1 transsynaptic spread from a genetically specified starter cell population. Resolving these two issues is therefore critical for expanding the application of this technique to reciprocally connected brain regions and refining its specificity as a circuit mapping tool. Future studies may seek to overcome these limitations by designing and exploring novel extracellular receptor and AAV capsid systems (Sun & Schaffer, 2018) that enable conditional uptake and transsynaptic transport of AAV in genetically specified neurons, but not neighboring cell-types or afferent terminals.

### Experimental Procedures

#### Animal preparation and stereotaxic surgery

All experimental procedures used in this study were approved by the Animal Care and Use Committee at the University of Southern California. Male and female Ai14 (Cre-dependent tdTomato reporter, Jackson Laboratories, stock #007914) and Frt-GFP (Flp-dependent GFP reporter, MMRRC, stock #32038) mice aged 2-6 months were used in this study. Mice were group housed in a light controlled (12 hr light: 12 hr dark cycle) environment with ad libitum access to food and water.

Stereotaxic injection of viruses was carried out as we previously described (Zingg et al., 2017; Ibrahim et al., 2016; Liang et al., 2015; Xiong et al. 2015). Mice were anesthetized initially in an induction chamber containing 5% isoflurane mixed with oxygen and then transferred to a stereotaxic frame equipped with a heating pad. Anesthesia was maintained throughout the procedure using continuous delivery of 2% isoflurane through a nose cone at a rate of 1.5 liters/min. The scalp was shaved and a small incision was made along the midline to expose the skull. After leveling the head relative to the stereotaxic frame, injection coordinates based on the Allen Reference Atlas (Dong, 2007) were used to mark the location on the skull directly above the target area and a small hole (0.5mm diameter) was drilled. Viruses were delivered through pulled glass micropipettes with a beveled tip (inner diameter of tip: ∼20 µm) using pressure injection via a micropump (World Precision Instruments). Total injection volumes ranged from 40 to 100 nL, at 15 nL/min. Following injection, the micropipette was left in place for approximately 5 mins to minimize diffusion of virus into the pipette path. After withdrawing the micropipette, the scalp was sutured closed and animals were administered ketofen (5mg/kg) to minimize inflammation and discomfort. Animals were recovered from anesthesia on a heating pad and then returned to their home cage.

For spinal injections of AAVretro-hSyn-GFP and AAVretro-hSyn-Cre, mice were anesthetized and positioned into a stereotaxic frame as described above. Cervical and lumbar injections were performed sequentially in the same procedure using aseptic technique. For each injection, a small patch of skin above the cervical or lumbar spinal region was shaved and a small incision was made to expose underlying muscle tissue. Muscle was blunt dissected and retracted to expose cervical (near C7-C8) or lumbar (near L3-L4) vertebrae and the spinal cord was stabilized using notched bars (Kopf, Model #987). Virus was injected unilaterally (200 nL total volume, 15 nL/min) between vertebral segments using a pulled glass micropipette with a beveled tip (inner diameter of tip: ∼20 µm) at a depth of 0.7 mm from the dorsal surface of the spinal cord. After withdrawing the micropipette, the skin was sutured closed and animals were recovered from anesthesia on a heating pad. To minimize inflammation and discomfort, animals were administered ketofen (5mg/kg) at the beginning of the surgical procedure and again every 24 hours for 4 days following surgery.

### Injection of viruses for anterograde transneuronal labeling

To compare the anterograde transneuronal transport properties of different viruses (Figure S1), self-complementary (sc) AAV1-hSyn-Cre (Vigene Biosciences, 2.8 × 10^13^ GC/mL), AAV1-hSyn-Flp (Vigene Biosciences, 5.5 × 10^13^ GC/mL), AAVretro-hSyn-Cre (Vigene Biosciences, 1.5 × 10^14^ GC/mL), AAVPHP.B-CMV-Cre (SignaGen, 2.3 × 10^13^ GC/mL), Adenovirus (Ad5-CMV-Cre, Kerafast, 3.0 × 10^12^ GC/mL), VSVG-pseudotyped Lentivirus (LV-CMV-Cre, Cellomics Tech, 1.0 × 10^8^ GC/mL), VSVG-pseudotyped Baculovirus (BAC-CMV-Cre, Uni. of Iowa, 3.7 × 10^10^ GC/mL), or G-deleted Rabies virus (RAV-Cre-GFP, Salk Institute, 8.6 × 10^8^ GC/mL) was injected into V1 (60 nL total volume; coordinates from bregma: anteroposterior −3.9 mm, mediolateral 2.6 mm, depth 0.5 mm) of Ai14 mice (for Cre-expressing viruses) or Frt-GFP mice (for AAV1-hSyn-Flp). Original titers were used for each virus injection. Animals were euthanized 4 weeks following injection and postsynaptic structures were examined for the presence of cell body labeling. The number of tdTomato+ cells in SC was quantified (see Imaging and Quantification section below) for each virus injection and plotted alongside previously reported data for additional viruses (Figure S1D, Zingg et al., 2017). Based on our results, we observed a nearly two-fold increase in the number of anterograde transsynaptically labeled cells when using AAV1-hSyn-Cre-WPRE as compared with scAAV1-hSyn-Cre or AAV1-hSyn-Flp, which both lack the woodchuck hepatitis post-transcriptional regulatory element (WPRE). This enhancer element can stabilize mRNA transcripts and increase the expression of AAV gene products by seven-fold (Loeb et al., 1999), however over-expression of Cre recombinase may become toxic to host cells at the injection site. To avoid this complication, all experiments in this study used scAAV1-hSyn-Cre and AAV1-hSyn-Flp, which do not exhibit any sign of toxicity at the injection site and still yield useful numbers of transsynaptically labeled cells (data not shown).

To test for the specificity of viral spread to different cell-types in the cerebellum, scAAV1-hSyn-Cre was injected into the PN (80 nL total volume; coordinates from bregma: anteroposterior −3.9 mm, mediolateral 0.4 mm, depth 5.5 mm) or IO (80 nL total volume; coordinates from bregma: anteroposterior −6.6 mm, mediolateral 0.4 mm, depth 5.5 mm) in Ai14 mice crossbred with GAD67-GFP mice (from Dr. Yuchio Yanagawa, Brain Science Institute, RIKEN, Japan). Animals were euthanized 2 weeks following injection and the cerebellar cortex was examined for the presence of tdTomato+ cell body labeling in the molecular, granule, and Purkinje cell layers.

To examine the efficiency of anterograde transsynaptic spread across different types of synapses, scAAV1-hSyn-Cre injections were targeted to inhibitory projection neurons or neuromodulatory cell populations in Ai14 mice. To test for viral spread through inhibitory synapses onto downstream inhibitory neurons, scAAV1-hSyn-Cre was injected into the Str (100 nL total volume; coordinates from bregma: anteroposterior +0.5 mm, mediolateral 2.3 mm, depth 3.0 mm) and the SNr was examined for cell body labeling (2 week post-injection survival time). To confirm the Str→SNr projection pathway is unidirectional, G-deleted Rabies-GFP (Salk Institute, 5.5 × 10^8^ GC/mL) was injected into the Str using the same coordinates as above and the SNc and SNr were examined for the presence of retrogradely labeled GFP+ cell bodies (50 nL injection, 1 week post-injection survival time). To test for viral spread through inhibitory synapses onto presumed excitatory neurons, SNr was injected with scAAV1-hSyn-Cre (50 nL total volume; coordinates from bregma: anteroposterior −3.3 mm, mediolateral 1.6 mm, depth 4.3 mm) and downstream neurons in VM were examined for the presence of tdTomato+ labeling (2 week post-injection survival time). To test for viral spread through neuromodulatory synapses, scAAV1-hSyn-Cre was injected (80 nL total volume, 2 week post-injection survival time) into the NDB (cholinergic neurons, coordinates from bregma: anteroposterior +0.5 mm, mediolateral 2.0 mm, depth 5.0 mm, 11° angle), DR (serotonergic neurons, coordinates from bregma: anteroposterior −4.5 mm, mediolateral 2.0 mm, depth 3.2 mm, 30° angle), or LC (noradrenergic neurons, coordinates from bregma: anteroposterior −5.4 mm, mediolateral 0.8 mm, depth 3.1 mm) of Ai14 mice and unidirectionally connected downstream targets were examined for cell body labeling (e.g. V1 for NDB and LC, or LGNd and ENTl for DR). To demonstrate the location and axonal targeting of V1-projecting cholinergic or LGNd-projecting serotonergic cell populations, AAV1-CAG-FLEX-GFP-WPRE (Addgene, 1.7 × 10^13^ GC/mL) was injected (60 nL total volume, 3 week post-injection survival time) into NDB of Chat-IRES-Cre mice (Jackson Laboratories, stock #006410) or DR of Pet1-Cre mice (Jackson Laboratories, stock #012712) using the same coordinates listed above.

To label different input-defined cell populations in the spinal cord, descending pathways in the brain were first identified by injecting AAVretro-hSyn-Cre (1.5 × 10^14^ GC/mL, 200 nL, Vigene Biosciences) and AAVretro-hSyn-GFP (1.7 × 10^14^ GC/mL, 200 nL, Vigene Biosciences) unilaterally into the left cervical (C7-C8) and lumbar (L3-L4) spinal cord, respectively. Animals were euthanized 3 weeks following injection and the entire brain was examined for retrogradely labeled GFP+ and/or tdTomato+ cell bodies. Several brain regions were then selected for injection of scAAV1-hSyn-Cre (100 nL total volume) in Ai14 mice using the following coordinates from bregma: MOp-ul (anteroposterior +0.7 mm, mediolateral 1.7 mm, depth 0.6 mm); MOp/MOp-ll (anteroposterior −0.8 mm, mediolateral 1.3 mm, depth 0.6 mm); SSp (anteroposterior −0.9 mm, mediolateral 1.7 mm, depth 0.6 mm); LHA (anteroposterior −1.5 mm, mediolateral 1.3 mm, depth 5.1 mm); RN (anteroposterior −3.5 mm, mediolateral 0.5 mm, depth 3.7 mm); and SUV (anteroposterior −5.8 mm, mediolateral 1.5 mm, depth 3.3 mm). Following a 2 week post-injection survival time, animals were euthanized and cervical, thoracic, and lumbar segments of the spinal cord were examined for tdTomato+ cell body labeling.

To reveal the topographical distribution of cells downstream of two different corticofugal pathways in the same brain, AAV1-hSyn-Flp was injected into MOp-ul (100 nL total volume, same coordinates as above) and scAAV1-hSyn-Cre was injected into MOp-ll (100 nL total volume, same coordinates as above) in Ai14-tdTomato × Frt-GFP mice. Following a 2 week post-injection survival time, downstream targets across the entire brain and spinal cord were examined for Flp+/GFP+ and Cre+/Tom+ cell body labeling.

To label the axonal output of PN neurons that specifically receive input from upper limb- or lower limb-related MOp, scAAV1-hSyn-Cre was injected into MOp-ul (80 nL total volume, same coordinates as above) or MOp-ll (same coordinates as above) and AAV1-CAG-Flex-GFP was injected into PN (80 nL total volume, coordinates from bregma: anteroposterior −3.9 mm, mediolateral 0.4 mm, depth 5.5 mm) in Ai14 mice. Following a 2 week post-injection survival time, animals were euthanized and the cerebellum was examined for GFP+ axons.

### Tetanus toxin expression

To explore underlying mechanisms of AAV transneuronal release, tetanus toxin light chain was expressed in V1 (same coordinates as described above) of Ai14 mice using injections of AAVDJ-CMV-TeNT-P2A-GFP (Stanford Viral Core, 5.7 × 10^12^ GC/mL, 100 nL total volume, n = 8 mice). As a control, a separate group of Ai14 mice were injected with AAV1-hSyn-GFP-WPRE (Addgene, titer reduced to 3.2 × 10^12^ GC/mL, 100 nL total volume, n = 8 mice). Following 2 weeks to allow for sufficient viral gene expression, scAAV1-hSyn-Cre was injected into the same location in V1 for each group (60 nL total volume). After an additional 2 weeks post-injection, mice were euthanized and SC was examined for the presence of tdTomato+ cell body labeling.

### Virus injections for sparse labeling of neurons

To achieve sparse labeling in a given Cre-expressing cell population, a co-injection strategy was used to obtain robust levels of YFP expression in only a few cells. Co-injections consisted of a 1:1 mixture of high titer AAVDJ-EF1a-fDIO-YFP-WPRE (UNC Vector core, final titer 1.2 × 10^13^ GC/mL) and low titer AAV1-EF1a-DIO-Flp-WPRE (Vigene Bioscience, diluted within a final titer range of 7.5 × 10^8^ – 10^10^ GC/mL). To establish a relationship between titer and the resulting number of labeled cells, AAV1-EF1a-DIO-Flp-WPRE was diluted to either 7.5 × 10^8^, 10^9^, or 10^10^ GC/mL and co-injected with AAVDJ-EF1a-fDIO-YFP-WPRE (50 nL total volume) in V1 of Ai14 × PV-Cre mice (Jackson Laboratories, stock # 017320, n = 4 mice for each titer). Following a 2 week post-injection survival time, animals were euthanized and V1 was examined for YFP+ cell labeling.

To achieve sparse labeling of both input- and genetically-defined cell populations, AAV1-EF1a-DIO-Flp-WPRE was injected at reduced titer (1.5 × 10^12^ GC/mL, 80 nL injection volume) into ACA (coordinates from bregma: anteroposterior +0.5 mm, mediolateral 0.3 mm, depth 0.9 mm) in Ai14 × Vglut2-Cre mice (Jackson Laboratories, stock #016963). This titer was chosen as previous results (Zingg et al., 2017) revealed the number of anterograde transsynaptically labeled cells in a given target region are reduced by 85-90% when injection titer is lowered from 10^13^ to 10^12^ GC/mL. A test injection of scAAV1-hSyn-Cre (80 nL volume, 1.5 × 10^13^ GC/mL) in ACA of Ai14 × GAD67-GFP mice revealed an average of about 8 GAD67-GFP+/Tom+ cells and 35 GAD67-GFP-/Tom+ cells (presumed excitatory) within a given 300 µm^3^ sample space of PAGdl. A reduced titer injection of AAV1-EF1a-DIO-Flp-WPRE (1.5 × 10^12^ GC/mL, 80 nL injection volume) in ACA of Vglut2-Cre mice would then be expected to tag ∼4 Vglut2-Cre+/Flp+ cells per 300 µm^3^ region of PAGdl. To robustly label these cells for morphological analysis, injections of AAVDJ-EF1a-fDIO-YFP-WPRE (30 nL total volume, 2.5 × 10^13^ GC/mL) were targeted to the PAGdl (coordinates relative to bregma: anteroposterior −4.0 mm, mediolateral 0.6 mm, depth 2.3 mm). Following a 2 week post-injection survival, animals were euthanized and PAGdl was examined for the presence of YFP+ cell bodies.

### Histology

Following desired post-injection survival time, animals were deeply anesthetized and transcardially perfused with 4% paraformaldehyde. Brains were extracted and post-fixed for 24 hours at 4°C in 4% paraformaldehyde and then sliced into 150 µm sections using a vibratome (Leica, VT1000s). The sections were serially mounted onto glass slides and coverslipped. For some experiments, a fluorescent Nissl stain was added (Neurotrace 640, ThermoFisher, N21483) to reveal cell body location and cytoarchitectural information.

### Imaging and quantification

All images were generated using a confocal microscope (Olympus FluoView FV1000). To quantify the total number of cell bodies labeled in SC following virus injection in V1 (Figure S1), serial sections across the entire structure were collected and examined. Regions with labeled cells were imaged at 10X magnification across the depth of the tissue (150 μm thickness, 15 μm z-stack interval). TdTomato+ cell bodies that co-localized with fluorescent Nissl stain were manually identified and counted.

To quantify cells labeled in the cerebellum following AAV1-Cre injection in PN or IO (Figure 1), coronal sections across the entire cerebellum were collected and the contralateral simple lobule (SIM) was selected for quantification as it exhibited strong and consistent labeling in all examined cases. For each animal, 10X magnification images were collected for 4 sections of SIM (from about −5.8 to −6.4 mm posterior to bregma) across the depth of the tissue (150 μm thickness, 15 μm z-stack interval). TdTomato+ neurons were quantified in the Purkinje cell layer and granule layer. To distinguish tdTomato+ granule cells from mossy fiber terminals, 40X z-stack images were collected throughout the granule layer and TdTomato+ cell bodies that co-localized with fluorescent Nissl stain were manually identified and counted. Cell counts for all 4 sections were totaled for each animal and plotted as mean ± SD (n = 4 mice for both PN and IO).

To provide an estimate of the efficiency of viral spread across different types of synapses (Figure 4), the percentage of tdTomato+ cells relative to Nissl+ cells was quantified within the axon terminal field in either SNr (for striatum injections) or VM (for SNr injections). For diffuse neuromodulatory output targeting entire structures or cortical regions, the percentage of tdTomato+ cells was quantified in relation to the total number of Nissl+ cells in the LGNd (for DR injections) or the total number of tdTomato+ cells relative to Nissl+ cells within a 500 × 500 µm sample space in V1 (for NDB and LC injections). 40X magnification images were used for quantification, and an average percentage was generated for each animal using at least 4 sample images in the target region (n = 4 mice for each pathway).

To quantify the number of sparsely labeled YFP+/Tomato+ PV cells in V1 following co-injections of AAV-fDIO-YFP and low titer AAV-DIO-Flp (Figure 8), serial sections through V1 were collected and imaged at 10X magnification across the depth of the tissue (150 μm thickness, 15 μm z-stack interval). The total number of YFP+/Tom+ cells were manually identified and counted for each of the titers tested (n = 4 animals each).

### Slice preparation and recording

To compare the rate of synaptic connectivity between tdTomato+, and neighboring non-labeled neurons in IC, a 1:1 mixture of scAAV1-hSyn-Cre and AAV1-EF1a-DIO-hChR2-YFP (Addgene, 1.6 × 10^13^ GC/mL) was injected into A1 of Ai14 mice (100 nL total volume, coordinates from bregma: anteroposterior −3.1 mm, mediolateral 4.5 mm, depth 0.7 mm). Following a 2 week post-injection survival time, acute brain slices containing IC were prepared. Following urethane anesthesia, the animal was decapitated and the brain was rapidly removed and immersed in an ice-cold dissection buffer (composition: 60 mM NaCl, 3mM KCl, 1.25 mM NaH2PO4, 25 mM NaHCO3, 115 mM sucrose, 10 mM glucose, 7 mM MgCl2, 0.5 mM CaCl2; saturated with 95% O2 and 5% CO2; pH= 7.4). Brain slices of 350 μm thickness containing IC were cut in a coronal plane using a vibrating microtome (Leica VT1000s). Slices were allowed to recover for 30 min in a submersion chamber filled with warmed (35 °C) ACSF and then to cool gradually to room temperature until recording. The spatial expression pattern of ChR2-EYFP in each slice was examined under a fluorescence microscope before recording. IC neurons were visualized with IR-DIC and fluorescence microscopy (Olympus BX51 WI) for specific targeting of both tdTomato+ neurons and nearby (within 150 µm) tdTomato-neurons surrounded by EYFP+ fluorescent fibers. Patch pipettes (Kimax) with ∼4-5 MΩ impedance were used for whole-cell recordings. Recording pipettes contained: 130 mM K-gluconate, 4 mM KCl, 2 mM NaCl, 10 mM HEPES, 0.2 mM EGTA, 4 mM ATP, 0.3 mM GTP, and 14 mM phosphocreatine (pH, 7.25; 290mOsm). Signals were recorded with an Axopatch 200B amplifier (Molecular Devices) under voltage clamp mode at a holding voltage of –70 mV for excitatory currents or 0 mV for inhibitory currents, filtered at 2 kHz and sampled at 10 kHz. 1 µM tetrodotoxin (TTX) and 1 mM 4-aminopyridine (4-AP) was added to the external solution for recording only monosynaptic responses (Petreanu et al. 2009) to blue light stimulation (3-10 ms pulse, 3 mW power, 10 trials, delivered via a mercury Arc lamp gated with an electronic shutter).

### Statistical Methods

All statistical analyses were performed using GraphPad Prism 6. Samples were first determined to have normal distribution using the Shapiro-Wilk test. To determine a significant association between anterograde transsynaptic labeling and synaptic connectivity (Figure 2D), Chi-square analysis was performed. To test for a significant difference in the number of cerebellar cell-types labeled (Figure 1I) or the number of cells found SC in tetanus toxin experiments (Figure 3D), unpaired Student’s t-test was used. Differences between data sets were considered significant if *p* < 0.05. Results for all histograms are expressed as mean ± SD.

