## Supplemental figure 1 and table 1 for "Anterograde Trans-Synaptic AAV Strategies for Probing Neural Circuitry"

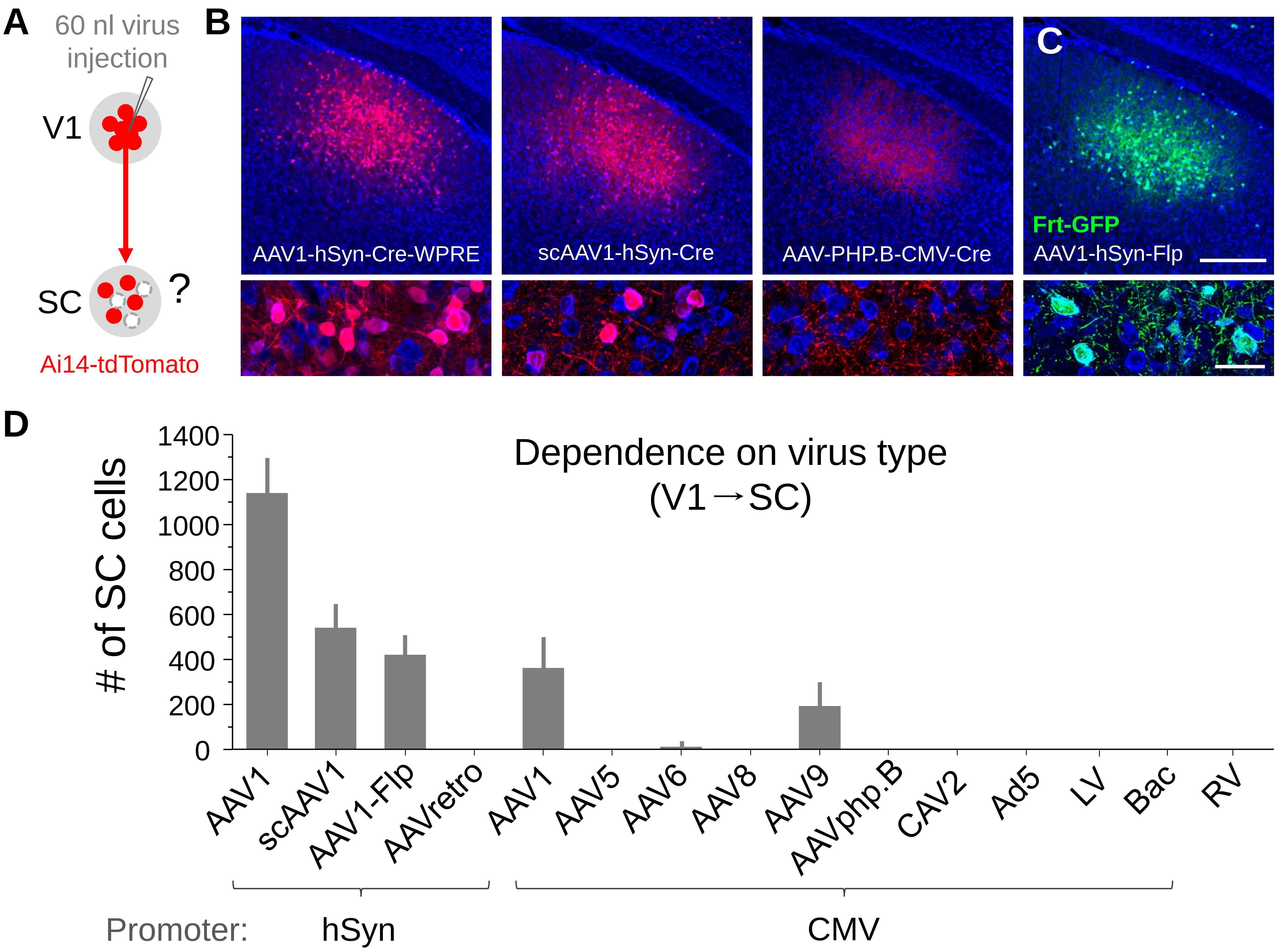


**Figure S1. Comparison of anterograde transsynaptic spread for different viruses.**

(A) Strategy for testing efficiency of transneuronal spread for different viruses. Injections of different Cre-expressing viruses (60 nL total volume) were targeted to primary visual cortex (V1) in Ai14-tdTomato Cre-reporter mice. Following a 4 week post-injection survival time, the superior colliculus (SC) was examined for Cre+/tdTomato+ cell bodies.

(B) Anterograde transsynaptic labeling of cell bodies (red) was observed in SC following V1 injections of AAV1-hSyn-Cre-WPRE (left) and self-complementary (sc) AAV1-hSyn-Cre (middle), but not AAV-PHP.B-CMV-Cre (right). Blue, fluorescent Nissl stain.

(C) Anterograde transsynaptic labeling using an alternate recombinase-reporter system. Flp+/GFP+ cells (green) were observed in SC following a 60 nl injection of AAV1-hSyn-Flp in V1 of Flp-dependent GFP reporter mice (Frt-GFP). Scale bars (B and C), 250 μm, top panels; 25 μm, bottom panels.

(D) Quantification of total number of cells labeled in SC for different Cre or Flp expressing viruses injected into V1 (4 week post-injection survival, 60 nL injection, n = 4 mice each). Error bar = SD.

**Table S1.** List of viruses used in this study.

| Virus | Titer | Source | Plasmid |
|---|---|---|---|
| AAV1-hSyn-Cre-WPRE | 2.5 x $10^{13}$ GC/mL | Addgene | 105553 |
| scAAV1-hSyn-Cre | 2.8 x $10^{13}$ GC/mL | Vigene Biosciences | |
| AAV1-hSyn-Flp | 5.5 x $10^{13}$ GC/mL | Vigene Biosciences | 51669 |
| AAVretro-hSyn-Cre | 1.5 x $10^{14}$ GC/mL | Vigene Biosciences | |
| AAV1-CMV-Cre | 2.7 x $10^{13}$ GC/mL | Addgene | 105537 |
| AAV5-CMV-Cre | 2.8 x $10^{13}$ GC/mL | Addgene | 105537 |
| AAV6-CMV-Cre | 3.5 x $10^{13}$ GC/mL | Addgene | 105537 |
| AAV8-CMV-Cre | 4.4 x $10^{13}$ GC/mL | Addgene | 105537 |
| AAV9-CMV-Cre | 1.6 x $10^{14}$ GC/mL | Addgene | 105537 |
| AAVPHP.B-CMV-Cre | 2.3 x $10^{13}$ GC/mL | SignaGen | |
| CAV2-CMV-Cre | 1.3 x $10^{12}$ GC/mL | Montpellier vector core | |
| Adenovirus (Ad5-CMV-Cre) | 3.0 x $10^{12}$ GC/mL | Kerafast | |
| Lentivirus (LV-CMV-Cre) | 1.0 x $10^{8}$ GC/mL | Cellomics Tech | |
| Baculovirus (BAC-CMV-Cre) | 3.7 x $10^{10}$ GC/mL | Uni. of Iowa | |
| G-deleted Rabies virus (RV-Cre-GFP) | 8.6 x $10^{8}$ GC/mL | Salk Institute | |
| G-deleted Rabies virus (RV-GFP) | 5.5 x $10^{8}$ GC/mL | Salk Institute | |
| AAV1-EF1a-DIO-Flp-WPRE | 1.5 x $10^{14}$ GC/mL | Vigene Biosciences | 87306 |
| AAVDJ-EF1a-fDIO-YFP-WPRE | 2.5 x $10^{13}$ GC/mL | UNC viral core | 55641 |
| AAV1-CAG-FLEX-GFP-WPRE | 1.7 x $10^{13}$ GC/mL | Addgene | 51502 |
| AAVretro-hSyn-GFP-WPRE | 1.7 x $10^{14}$ GC/mL | Vigene Biosciences | 105539 |
| AAV1-hSyn-GFP-WPRE | 3.2 x $10^{13}$ GC/mL | Addgene | 105539 |
| AAV1-EF1a-DIO-hChR2-eYFP-WPRE | 1.6 x $10^{13}$ GC/mL | Addgene | 20298 |
| AAVDJ-CMV-TeNT-P2A-GFP | 5.7 x $10^{12}$ GC/mL | Stanford viral core | |
| AAVretro-EF1a-Cre-WPRE | 2.3 x $10^{13}$ GC/mL | Salk Institute | 55636 |
